# An evaluation of pool-sequencing transcriptome-based exon capture for population genomics in non-model species

**DOI:** 10.1101/583534

**Authors:** Emeline Deleury, Thomas Guillemaud, Aurélie Blin, Eric Lombaert

## Abstract

Exon capture coupled to high-throughput sequencing constitutes a cost-effective technical solution for addressing specific questions in evolutionary biology by focusing on expressed regions of the genome preferentially targeted by selection. Transcriptome-based capture, a process that can be used to capture the exons of non-model species, is use in phylogenomics. However, its use in population genomics remains rare due to the high costs of sequencing large numbers of indexed individuals across multiple populations. We evaluated the feasibility of combining transcriptome-based capture and the pooling of tissues from numerous individuals for DNA extraction as a cost-effective, generic and robust approach to estimating the variant allele frequencies of any species at the population level. We designed capture probes for ∼5 Mb of chosen *de novo* transcripts from the Asian ladybird *Harmonia axyridis* (5,717 transcripts). We called ∼300,000 bi-allelic SNPs for a pool of 36 non-indexed individuals. Capture efficiency was high, and pool-seq was as effective and accurate as individual-seq for detecting variants and estimating allele frequencies. Finally, we also evaluated an approach for simplifying bioinformatic analyses by mapping genomic reads directly to targeted transcript sequences to obtain coding variants. This approach is effective and does not affect the estimation of SNP allele frequencies, except for a small bias close to some exon ends. We demonstrate that this approach can also be used to predict the intron-exon boundaries of targeted *de novo* transcripts, making it possible to abolish genotyping biases near exon ends.

## Introduction

One of the core challenges in population genomics and phylogenetics is to obtain genotype calls for a reliable set of orthologous loci from a sufficient number of individuals across multiple populations and from species spanning a range of divergences, respectively. The targeted resequencing of a consistent subset of genomic regions is a reproducible and cost-effective technical solution, with much lighter data handling than for whole-genome sequencing. Hybridization capture is one of these reduced representation methods that allows the enrichment of a preselected set of hundreds to thousands of genes or DNA fragments from the genomic DNA (Hodges *et al*., 2007). The most common application of hybridization sequence capture is exon capture, in which only coding regions of the genome (and their flanking regions) are sequenced (Warr *et al*., 2015). This approach provides focused information about gene function and adaptation, and, unlike transcriptome sequencing, is not biased by variations of gene expression.

Exome capture requires a knowledge of the exon sequences of the species studied, for the design of appropriate oligonucleotide probes. Consequently, its use has been largely limited to model species (i.e. species for which a reference genome is available). Absolute probe specificity is not required, however, and given the tendency of functional elements to be conserved even after millions of years of divergence, exons can be captured from a non-model species (i.e. a species without a published genome sequence) with probes designed from the sequence of a related model species (e.g. Cosart *et al*., 2011; Förster *et al*., 2018). This approach has been used for the successful resolution of species phylogenies (e.g. Ilves & López-Fernández, 2014; Bossert & Danforth, 2018; Ilves *et al*., 2018). It is also an effective method for capturing and enriching degraded and contaminated ancient DNA from extinct groups in paleontology (e.g. Castellano *et al*., 2014).

An alternative approach for non-model species involves designing DNA capture probes on the basis of sequence subset obtained from a *de novo* assembled transcriptome for the species studied or a related species (e.g. Bi *et al*., 2012; Neves *et al*., 2013). One of the limitations of this approach is that the intron-exon structure is unknown, and probes based on transcript sequences spanning two consecutive exons therefore hybridize only partially to genomic DNA, reducing capture efficiency (e.g. Neves *et al*., 2013). One way of addressing this issue is to choose only targets which have orthologous genes in closely related model species to predict intron positions (Bi *et al*., 2012; Stephens *et al*., 2015; Bragg *et al*., 2016). However, the use of data from closely related model species biases target choice toward the retention of genes displaying the highest degree of conservation throughout evolution, which is potentially problematic in population genomics studies. In addition, even when orthologous genes are targeted, transcriptome-based exon capture studies have still been found to have a lower capture efficiency close to intron-exon boundaries, because it is not possible to design capture probes spanning exons and their (unknown) non-coding flanking regions (Bi *et al*., 2012; Neves *et al*., 2013; Portik *et al*., 2016).

Captured fragments are usually longer than probes. Transcriptome-based exon capture therefore provides opportunities to capture the flanking regions of the exons, thereby expanding gene models beyond the available coding sequence (Bi *et al*., 2012; Neves *et al*., 2013; Stephens *et al*., 2015; Bragg *et al*., 2016; Portik *et al*., 2016). This requires the construction, in a fragmented way (e.g. long introns not fully captured), of the genomic sequences corresponding to the targeted transcripts (e.g. Neves *et al*., 2013). This assembly (generally referred to as an in-target assembly) is used as a reference for the mapping of genomic reads or for phylogeny construction. This assembly is obtained by cleaning the raw genomic sequences and assembling them with various k-mer combinations. The genomic contigs obtained are then compared with the target sequences for ordering and scaffolding with each other when they correspond to the same targets. Such a complex approach is necessary for phylogeny construction when probes are used to capture orthologous targets from related species, but may be dispensable in the context of population genomics. Indeed, several studies have shown that the assembly process does not completely rebuild all targets (e.g. Bragg *et al*., 2016; Portik *et al*., 2016), and this may decrease the likelihood of detecting variants. Bragg et al. (2016) tried to attenuate this problem by mapping the genomic reads of the species used for probe design directly onto the transcript target sequences, thereby making it possible to cover exons that were not successfully assembled but were nevertheless captured and sequenced.

Transcriptome-based exon capture has proved effective for the generation of thousands of highly informative markers. Most studies to date have aimed to resolve the phylogenetic relationships between various set of related species (Nicholls *et al*., 2015; Stephens *et al*., 2015; Bragg *et al*., 2016; Heyduk *et al*., 2016; Ilves *et al*., 2018), and applications in population genomics remain rare (but see Bi *et al*., 2013; Rellstab *et al*., 2019). This is probably because population studies traditionally analyze large numbers of individuals from multiple populations of non-model species, which is complex, time-consuming and costly, even if sequencing capacity and costs are optimized by indexing individuals for multiplexing (Shearer *et al*., 2012). However, many research questions in population genomics can be addressed by simply using the allele frequencies calculated at population level (e.g. Gutenkunst *et al*., 2009). In this context, pooling individuals within the population before the extraction of DNA may be an efficient, cheap and time-saving strategy. Pool-sequencing has been shown to estimate allele frequencies within populations as accurately as individual genotyping, but with much less effort for library construction and sequencing (e.g. Gautier *et al*., 2013; Schlötterer *et al*., 2014). Allele frequencies have been estimated by a combination of targeted capture methods and pool-sequencing strategies only in humans (Bansal *et al*., 2011; Day-Williams *et al*., 2011; Ramos *et al*., 2012; Ryu *et al*., 2018) and, very recently, in pine trees (Rellstab *et al*., 2019).

We propose here an efficient way of quantifying the single-nucleotide polymorphism (SNP) of exomes at the population level in non-model species for which no genome sequence is available. This approach is based on a combination of transcriptome-based exon capture and the pooling of tissues from numerous individuals before DNA extraction. We report the development of capture probes on a subset of the exome of the Asian ladybird *Harmonia axyridis* from its unannotated available *de novo* transcriptome (Vilcinskas *et al*., 2013; Vogel *et al*., 2017), without the use of the available draft genome (HaxR v1.0; Gautier *et al*., 2018) and, thus, in the absence of known gene structure, as in most non-model species. We used these probes for two target enrichment experiments with the same population sample, the first based on capture with indexed multiplexed individuals, and the second based on capture with a pool of non-indexed individuals. We evaluated the efficiency of capture and the accuracy of the estimates of allele frequencies obtained by pool sequencing in this context. We also evaluated a rapid approach for the processing of read data without their assembly, based on the mapping of reads directly to targeted *de novo* transcript sequences for the detection of variant loci (Bragg *et al*., 2016; Rellstab *et al*., 2019). This direct mapping also makes it possible to effectively predict the locations of the intron-exon boundaries of target sequences.

## Methods

The overall analysis workflow is summarized in Figure 1.

**Figure 1:**
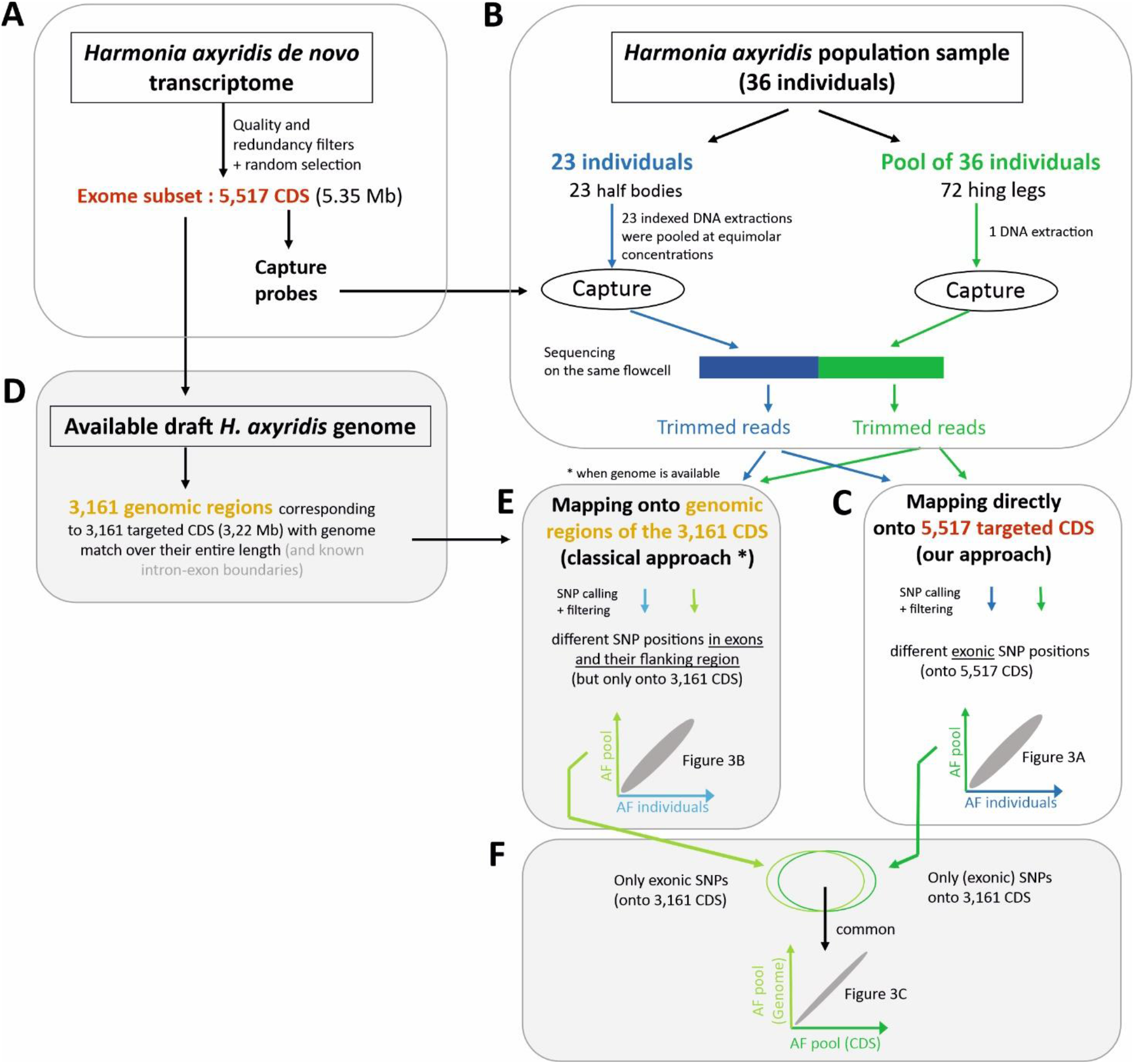
Overview the analysis workflow (see text for details).

### Design of capture probes for target enrichment

Using an already available *de novo* transcriptome obtained by RNA-Seq on various tissues and conditions/stages of *H. axyridis* (Vilcinskas *et al*., 2013; Vogel *et al*., 2017), we sought putative peptide-coding sequences with FRAMEDP (v1.2.2, Gouzy *et al*., 2009), using BLASTX results (e-value ≤ 1e^-7^) of transcripts against the insect proteomes of *Tribolium castaneum, Anoplophora glabripennis, Dendroctonus ponderosae, Drosophila melanogaster* and the SwissProt database (v2016-02-17) for training. We have chosen to work on an exome subset, particularly with the future goal of working simultaneously on a large number of population samples. To maximize the effectiveness of our probes, target sequences were selected as follows. The corresponding partial or complete coding sequences (CDS) were filtered to eliminate (i) *Wolbachia* and other putative endosymbiont sequences, (ii) CDS with more than 1% missing data (*Ns*) or with more than four consecutive *Ns*, (iii) CDS with a GC% below 25 or above 75 and (iv) CDS with lengths below 400 bp (to avoid the capture of too great a proportion of flanking regions and to allow better tiling) or above 3500 bp (to make it possible to target larger numbers of different genes). We identified 28,326 putative CDS, of which 12,739 were retained after the application of the filters. We then chose a subset of 5.5 Mb at random: the index (1 to 12,739) of the CDS was drawn from a discrete uniform distribution U[1;12,739] without replacement until the desired subset exome size is reached.

The probes based on the selected CDS were designed and produced by NimbleGen. The repetitiveness of the probe sequences was assessed on the basis of the frequency of 15-mers in the genomes of the coleopterans *T. castaneum* and *A. glabripennis*. Probes with more than one close match, defined as no more than five single-base insertions, deletions or substitutions with the SSAHA algorithm (Ning *et al*., 2001), in the *H. axyridis de novo* transcriptome or in the draft genome of *A. glabripennis* were discarded. Probes matching to sequences in the *H. axyridis* or *T. castaneum* mitochondrial genome were also discarded. The few residual *Ns* in target sequences were replaced with a random nucleotide to allow tile probing across a single unknown base. The final probe set contained 6,400 regions of overlapping probes, corresponding to a targeted exome size of 5,347,461 bases distributed over 5,717 of our selected CDS (Fig. 1A; Zenodo https://doi.org/10.5281/zenodo.2598388). This final probe set was used for the preparation of a SeqCap EZ Developer assay by Roche NimbleGen Technologies, which synthesized the probes as biotinylated DNA oligomers. One capture reaction contained 2,100,000 overlapping probes of 50 to 99 bp in length, with a mean length of 74.71 ± 4.92 bp.

### Individual and pooled genomic DNA library preparation

Thirty-six individuals of *H. axyridis* from a wild population in China (Beijing; 40.057128°N, 116.53989°E) were collected in October 2015, immediately placed in RNAlater solution and stored at −20°C. Based on the PIFs (“Pool Individual Frequency SNP”) tools proposed by Gautier *et al*. (2013), we would theoretically expect a similar precision of allele frequency estimation for the sequencing of 23 individually indexed individuals, each at a sequencing depth of 80X, or a single pool of 36 individuals at 125X, assuming an experimental error (i.e. ε, the coefficient of variation of the contribution of each individual to the set of sequence reads obtained from the DNA pool) of 50%. We thus extracted DNA (i) independently from 23 individuals, using half the body (without the hind legs), and (ii) from a pool of 36 individuals, including the 23 individuals used for individual sequencing, using two hind legs per individual (Fig. 1B). All DNA extractions were performed with the Qiagen DNeasy Blood & Tissue kit, in accordance with the manufacturer’s recommendations.

Genomic libraries were prepared with NimbleGen SeqCap EZ Library (v5.0). Briefly, for each of the 24 samples (1 pool and 23 individuals), DNA (2 µg in 100 µl) was mechanically sheared to a mean fragment length of 200 bp with a Covaris S2 E210 device (6×30s). Fragments were end-repaired, A-tailed and indexed (one index per sample) with KAPA Library Preparation Kit Illumina platforms. After the ligation of Illumina adaptors and indexes, only fragments with a length of 250 to 450 bp were retained. PCR was performed (7 cycles) with standard Illumina paired-end primers and the amplicons were then purified with AMPure XP beads (Beckman). The length, quality and concentration of the prepared DNA fragments were checked on a BioAnalyzer (Agilent High Sensitivity DNA Assay) and a Qubit.

### Sequence capture hybridization and sequencing

The 23 indexed individuals were pooled at equimolar concentrations before hybridization. For each capture (one with the pool and one with the indexed individuals), a total of 1 µg of amplified DNA was used for exome enrichments, with the SeqCap EZ Developer probes described above, strictly in accordance with the NimbleGen SeqCap EZ Library protocol (v5.0). For each capture, two parallel post-capture PCRs (14 cycles) were performed on elution solution, and the PCR products were then merged before purification with AMPure XP beads. The length, quality and concentration of the final DNA fragments were checked on a BioAnalyzer (Agilent High Sensitivity DNA Assay) and a Qubit. Each capture was sequenced on a half lane of an Illumina HiSeq3000 sequencer (on the same lane; Fig. 1B), in accordance with the manufacturer’s instructions, in paired-end mode, for 150 cycles. The data were then demultiplexed and exported as FastQ files, and the 24 libraries were processed independently.

### SNP calling by the mapping of reads to the CDS

Sequence quality was checked with FastQC (https://www.bioinformatics.babraham.ac.uk/projects/fastqc/). Adapters were removed and low-quality base pairs were trimmed with Trimmomatic (v0.35, Bolger *et al*., 2014), with the following parameters: ILLUMINACLIP:TruSeq-file.fa:2:30:10 LEADING:30 TRAILING:30 SLIDINGWINDOW:10:30 MINLEN:75, and FASTX (v0.0.14; http://hannonlab.cshl.edu/fastx_toolkit/) to remove the first five bases.

We chose to align the reads directly with the targeted CDS sequences, because target enrichment was performed on the same species as probe design, as suggested by Bragg *et al*. (2016; see also Rellstab *et al*., 2019) (Fig. 1C). Doing so, we avoid the complex and time-consuming steps of read assembly and scaffolding/ordering the genomic contigs obtained to generate the in-target assemblies for read mapping (see introduction). For each library, cleaned reads were aligned with the 5,717 targeted CDS, with BOWTIE2 (v2.2.5; Langmead & Salzberg, 2012), using the default parameters and *local* option to allow for genomic reads not aligning with the targeted CDS sequence over their entire length, whilst ensuring a minimum alignment length of 20 bp. Reads were aligned as unpaired, to allow for the mapping of orphan reads. Reads with multiple alignments, and those for which the mate-pair was aligned to a different target were discarded after mapping. The aligned data in SAM format were converted into BAM format and sorted with SAMtools (v0.1.19). Duplicate reads were estimated with picard tools (v1.113) MarkDuplicates, using the default settings. Coverage at each target base was calculated with mosdepth (v0.2.3; Pedersen & Quinlan, 2018).

Variant calling was performed with read alignments of more than 40 bases in length containing fewer than 10 deletions, as deduced from the *cigar* format. SNP sites were called separately for each of the 24 libraries with the SAMtools *mpileup* command, using option *-B* followed by the varscan2 (v2.3; Koboldt *et al*., 2012) *mpileup2snp* command with a minimum base quality of 35 to count a read at a given position, and a *p*-value threshold of 0.01. For the other parameters, we considered only positions with a minimum sequencing depth of 40 reads for each individual, and 250 reads for the pool (i.e. for these selected depths, the same exome size was analyzed: 90.3% and 93% of target bases were supported by at least 40 and 250 reads for the individual and pooled exomes, respectively; see Fig. S1). We also only included a variant if it was supported by 15 reads per individual (to take into account the variance between both haplotypes sequencing depths), or by three reads for the single pool (given the pool’s minimum sequencing depth of 250X, we expected to have ∼3 reads per haplotype if the 72 haplotypes in the pool were homogeneously captured). Moreover, to account for the variation in total read sequencing depth across positions, we required a minimum variant allele frequency threshold of 0.25 for each individual and 0.01 for the pool. For individual libraries, an individual was considered to be heterozygous if the variant allele frequency f(v) was between 0.25 and 0.75.

All 409,328 different SNP positions independently found in the 24 libraries were then genotyped for each library with varscan2 *mpileup2cns*, using the same parameters as described above, but without imposing a minimum number of reads with variant for individuals. Several filters were then applied to all SNP positions, using either in-house PERL or R scripts (R Development Core Team, 2015). We first retained only bi-allelic positions. In order to properly estimate allelic frequencies with individuals, we retained only positions genotyped for at least 20 individuals. Finally, we discarded SNPs with high levels of sequencing depth in the pool, which were likely to represent repetitive regions or paralogs of the genome: maximum pool sequencing depth was set at 6,960X, as calculated with the formula proposed by Li et al. (2014): *c*+4*√*c*, where *c* is the mean sequencing depth in the pool (*c*=6,634.8X). We retained 300,036 bi-allelic SNPs (73.3%) after the application of all these filters (Fig. 1C).

At each site, and for each capture, variant allele frequency at population level was estimated as the number of reads with the variant allele divided by total sequencing depth. Individual genotypes were used to calculate exact *p*-values for Hardy-Weinberg equilibrium tests with the “HardyWeinberg” R package (Graffelman, 2015), with the correction of significance level by the false discovery rate procedure (Benjamini & Hochberg, 1995). Finally, SNPs were annotated and categorized with the SnpEff program (v4.3t; Cingolani *et al*., 2012).

### Evaluation of the approach based on read mapping directly onto CDS

Approaches based on the direct mapping of reads onto CDS have already been used to detect SNPs (e.g. Rellstab *et al*., 2019), but they have never been evaluated. We used an *H. axyridis* draft genome sequence (HaxR v1.0; Gautier *et al*., 2018) to evaluate this approach and that proposed below for the prediction of intron-exon boundaries (IEBs), but only for a subset of our targets, i.e. those with a genomic match (Fig. 1D). We selected CDS targets with a genomic match, as proposed by Neves *et al*. (2013), with some adjustments. Briefly, CDS target sequences were first aligned with genomic scaffolds with BLASTN (e-value cutoff of e^-20^ and identity cutoff of 90%), and the best hit was retained for each target. The target sequences were then realigned with the corresponding genomic scaffold, with both GMAP (version 2015-12-31; Wu & Watanabe, 2005) and EXONERATE *est2genome* (v2.2.0; http://www.genome.iastate.edu/bioinfo/resources/manuals/exonerate/). We isolated targets (i) with genome matches over their entire length, (ii) for which intron/exon structure predictions were the same for both alignment programs, (iii) which were covered by capture probes over their entire length and (iv) for which we had access to at least 200 bp upstream and downstream from the targeted CDS sequences. With these filters, we retained 3,161 CDS targets in total (3,220,718 bp, i.e. 60.23% of our targeted exome subset). These 3,161 full or partial CDS corresponded to 11,754 exons, with a mean of 3.72 exons per target (range: 1 to 17 exons) and a mean exon length of 274 bp (range: 12 to 3,417 bp). These 11,754 exons included 6,295 (53.6%) that were unambiguously complete, because they were surrounded by two others, and 8,593 reliable IEBs in total were identified between consecutive exons.

All genomic regions corresponding to these 3,161 CDS targets were then isolated, including 200 bp upstream and downstream from the target sequences (Fig. 1D). Independently for each library, all cleaned reads were aligned with these genomic regions, with BOWTIE2 (v2.2.5), using the default parameters and the *end-to-end* option (i.e. reads must be aligned along their full length). Reads with multiple alignments, and reads for which the mate-pair was aligned to another target were discarded after mapping. Coverage calculation, SNP calling and variant allele frequency estimation were performed as described above (Fig. 1E). We established the correspondence between the position of a base in the genomic region and its position in the CDS sequence, using the *vulgar* format of EXONERATE output (Fig. 1F). SNP positions for which there was a deletion or an insertion between the CDS sequence and its associated genomic sequence (i.e. 559 positions) were excluded from the comparison.

In order to compare quantitatively the allelic frequencies obtained with the different approaches presented here (i.e. pool vs. individuals; CDS mapping vs genome mapping), we calculated in each case the Pearson correlation coefficient and the Bland-Altman average bias with its standard deviation which provide information upon agreement between measurements (Altman & Bland, 1983).

### Intron-exon boundary detection

Our approach is based solely on the direct mapping of genomic reads to the CDS sequences, by allowing reads to map over less than their full length (i.e. local alignment). Thus, a genomic read with both part of an exon and its flanking sequences (intron or UTR) will start mapping to the corresponding exon in the CDS sequence, and stop mapping at the exon end (Conklin *et al*., 2013). This will generate a signal of the intron-exon boundary in the number of reads that either begin or end their local alignment at that position (hereafter referred to as *nbBE*). Along a covered exon, *nbBE* should be much lower than the sequencing depth rate, except at the end of the exon, where we should observe a large increase in *nbBE* to a value theoretically equal to the sequencing depth (Fig. S2A). This strong increase in *nbBE* should occur either at the base at the exact end of the exon, or a few bases away if the start of the intron sequence is, by chance, highly similar to the start of the next exon sequence on the CDS (Fig. S2B). In this second case, it creates an abnormally inflated estimate of sequencing depth locally for a few bases at the boundary of the following exon (Fig. S2B and Fig. S3 for an example). At an IEB (i.e. a junction of two exon ends in the CDS), depending on whether a signal offset occurs for one or both exon ends, the size of the area with abnormally inflated local sequencing depth is variable, but generally less than 10 bp. In addition, the presence of a SNP at some bases at the end of the exon may prevent reads with the variant from aligning up to the IEB position, potentially generating another signal in addition to those of the two exon ends of the IEB. Combinations of these different scenarios may also occur.

We developed and tested a method for predicting IEBs on the basis of the increase in *nbBE*, by focusing on the 3,161 targeted transcripts for which genomic sequences were available (Fig. 1D). We used the mapping output BAM file obtained with the pool library (Fig. 1C). At each position in the transcript, *nbBE* was calculated from the start read alignment position on the transcript and the *cigar* code, both present in the BAM file. The value of the *nbBE* parameter at each position is a function of sequencing depth (and read length). We took the possible effect of sequencing depth heterogeneity on *nbBE* into account by calculating the median *nbBE* over a window of 11 bases centered on the position concerned. A signal (i.e. a predicted IEB), was detected at a position if *nbBE* was *X* times larger than the median, *X* being a tool parameter defined by the user. For the definition of *X*, it may be necessary to identify a subset of transcripts with orthologs in a close model species, and then to determine which value of *X* provides the best prediction of IEBs for this subset. Signals at the exact ends of the CDS were not considered. We took the possible signal deviation (i.e. a signal that is slightly deported with respect to the IEB that generates it; Fig. S2) into account, by extending signal detection to the three bases upstream, and three bases downstream from a position with measured signal (default value for parameter *n*=3). A measured signal therefore predicts an IEB in a window of seven bases in total. An IEB, composed of two exon ends, theoretically generates two close signals with overlapping windows. If these two signals are found at the exact positions of the exon ends (no deviation), then the predicted region measures 8 bp (Fig. S2A). If these two signals overlap (e.g. because of the deviation of one of the two signals) then the predicted region for the IEB is 7 bp long. It can be less than 7 bp if the signal is detected close to non-covered regions or regions with measured signals close to the end of the CDS for which signal extension is not possible. If these signals deviate by up to six bases from each other (i.e. cumulative deviation of eight bases, the deviation being in opposite directions for the two signals) then the predicted region measures 14 bp (Fig. S2). More rarely, if a third signal is detected in the same region, it increases the length of the region by a few bases. If the two signals deviate by more than six bases from each other, then the tool predicts two signal windows separated by a few bases for the IEB concerned. Thus, each transcript was divided into short regions with prediction signals (+) alternating with regions without prediction signals (-). Each region was then assessed as follows: true positive (TP) if the + region contained an IEB; false positive (FP) if the + region did not contain an IEB; true negative (TN) if the - region did not contain an IEB; and false negative (FN) if the - region contained an IEB. We evaluated the performance of the method, by calculating the percentage of true IEBs correctly predicted (i.e. sensitivity (SN)) as TP/(TP+FN) and the percentage of correct predictions (i.e. specificity (SP)) as TP/(TP+FP) (Burset & Guigo, 1996). For a CDS with no introns (862, i.e. 27.3% of the 3,161 CDS), SN cannot be evaluated; SP was 100 if no IEB was predicted (good prediction) and 0 otherwise. Scripts are available from https://github.com/edeleury/IEB-finder.

## Results

### Capture data and filtration

In the two target enrichment experiments, we obtained 186,998,614 and 185,457,659 *H. axyridis* raw sequence read pairs for the pool library (SRA ERX3237193) and the 23 indexed individuals (SRA ERX3237194 to ERX3237216), respectively. For each indexed individual library, we obtained a mean of 8,063,376 raw read pairs (range: 6,912,561 to 9,176,292). After cleaning, 91.2% of the raw read sequences were retained (see detailed information for all libraries in Table S1).

### Capture efficiency, sequencing depth and duplicates

Rather than assembling the cleaned reads, we chose to map them directly onto the target CDS sequences used for probe design (Fig. 1C). Mean capture specificity, defined as the percentage of cleaned reads mapping correctly (i.e. meeting our mapping parameters) onto the 5,717 targeted CDS, was 83.2% (range over libraries: 80.6%-89.2%; Table S2). Aligned bases accounted for 74.6% (range over all libraries: 72.6%-75.7%) of all cleaned read bases. The global median read alignment length was 144 bp (range over libraries: 143-144 bp, see Table S2), with 62.7% of the mapped reads (range over all libraries: 60.9%-64.3%) mapped over their entire length. A substantial proportion of the mapped reads (22.2%; range over all libraries: 20%-25.9%) were ‘orphaned’ mates (i.e. only one of the two mates aligned to the target sequences). This percentage strongly suggests that flanking regions of targeted regions were also sequenced with our capture procedure (Yi *et al*., 2010; Neves *et al*., 2013), although quantification was not possible with this direct mapping approach.

The mean percentage of the 5,347,461 targeted bases (5,717 target CDS) covered by at least one read (i.e. the capture sensitivity) was 93.5% (range over all libraries: 93.4%-93.9%, see Fig. S1 and Table S3). Overall, 93% of the targeted bases were covered in all 24 libraries, whereas 6% were covered by no read in any of the libraries (Table 1). We identified 491 full targets (292,327 bp) with no read aligned in the 24 libraries. For these uncovered targets, 487 (99.2%) had no match to the *H. axyridis* draft genome either, whereas, for the 5,226 covered targeted CDS, only 74 (1.4%) had no match to the genome (here partial alignments of targets with genome were also considered).

**Table 1:** Reproducibility of capture sensitivity on the 24 libraries. 100% means that all 24 samples had at least one read aligned with the target, whereas 0 % level indicates that no reads aligned with the target.

| Target | Target size | Percentage of samples with target sequence |  |
| --- | --- | --- | --- |
|  |  | 100% | 0% |
| CDS level | 5 717 | 5 226 (91.4%) | 491 (8.6%) |
| Base level | 5 347 461 | 4 967 690 (92.9%) | 323 418 (6.0%) |

Considering all targeted bases, the median and mean base sequencing depth for individual libraries were 142X and 283X for the “23 individuals” libraries, and 3,361X and 6,635X for the pool libraries (see Table S3), respectively. A small number of targeted bases had very high sequencing depth, but around 90% had a depth <300X for individuals or <7,000X for the pool (Table S3). For individuals, 90.3% of the targeted bases on average had a mean sequencing depth greater than 40X. For the pool, 93% had a depth greater than 250X (Fig. S1).

Base sequencing depth varied along the targeted CDS (Fig. S4A). Using the known complete exons (≥ 20 bp) identified on targeted CDS with a match over their entire length with the *H. axyridis* draft genome, we confirmed that mean sequencing depth increased steadily with exon size (Fig. S4B), and that base sequencing depth decreased from the middle of an exon to its ends (Fig. S4C). Alignments of less than 20 bp between the reads and the CDS sequence were not permitted with this mapping procedure, so sequencing depth was probably underestimated at exon ends and for short exons. For all the targeted CDS considered together, mean sequencing depth differed between targeted CDS but was highly reproducible between libraries (Fig. S5): the Pearson correlation coefficient of the target sequencing depth between individual libraries ranged from 0.96 to 0.99, with a mean of 0.99, and the Pearson coefficient of correlation between the two capture processes (i.e. between pool and individuals) was 0.999.

For the 23 individual libraries, a mean of 66.1% of reads were scored as duplicates (range over libraries: 63.6%-68.9%; Table S4). This high proportion was due to the very strong sequencing effort regarding the target region size. Indeed, pooling the reads of the 23 individual libraries increased the proportion of duplicates detected to >96%, as for the pool (Table S4), suggesting that the reads scored as duplicates in our experiment probably mostly come from different individuals. We did therefore not discard duplicated reads before SNP calling, so as to avoid artificially biasing the estimation of allele frequencies.

### SNP calling and accuracy of allele frequency estimation

SNP calling performed independently for the individuals and for the pool, following direct mapping onto the targeted CDS sequences, led to the detection of 302,292 and 364,144 different putative exonic SNPs, respectively. Among the total 409,328 SNPs, 257,108 positions (62.8%) were detected in both experiments, 45,184 (11%) only in individuals and 107,036 (26.2%) only in the pool. As expected (because there were 13 more individuals in the pool), the number of SNPs called for the pool was larger than that for the 23 individuals, with 26.2% of SNPs called for the pool only. The vast majority (95.6%) of the SNPs called only for individuals had allele frequencies below 0.03.

All positions were genotyped in all libraries. After filtering, we finally selected a total of 300,036 exonic bi-allelic SNPs within 4,741 targeted CDS (Fig. 1C). Minor allele frequencies (MAFs) were below 0.03 for 59.2% of these SNPs (Fig. 2); 92.75% were polymorphic in both the pool and the individuals, 7.22% were polymorphic only in the pool, and 0.03% were polymorphic only in the individuals. Only 1,007 of the 300,036 SNPs (i.e. 0.34%) displayed significant deviation from Hardy-Weinberg equilibrium, as calculated with the individual genotypes. This number was more than three times higher (i.e. 3,331 loci) before the removal of loci on the basis of the maximum sequencing depth filter applied to the pool.

**Figure 2:**
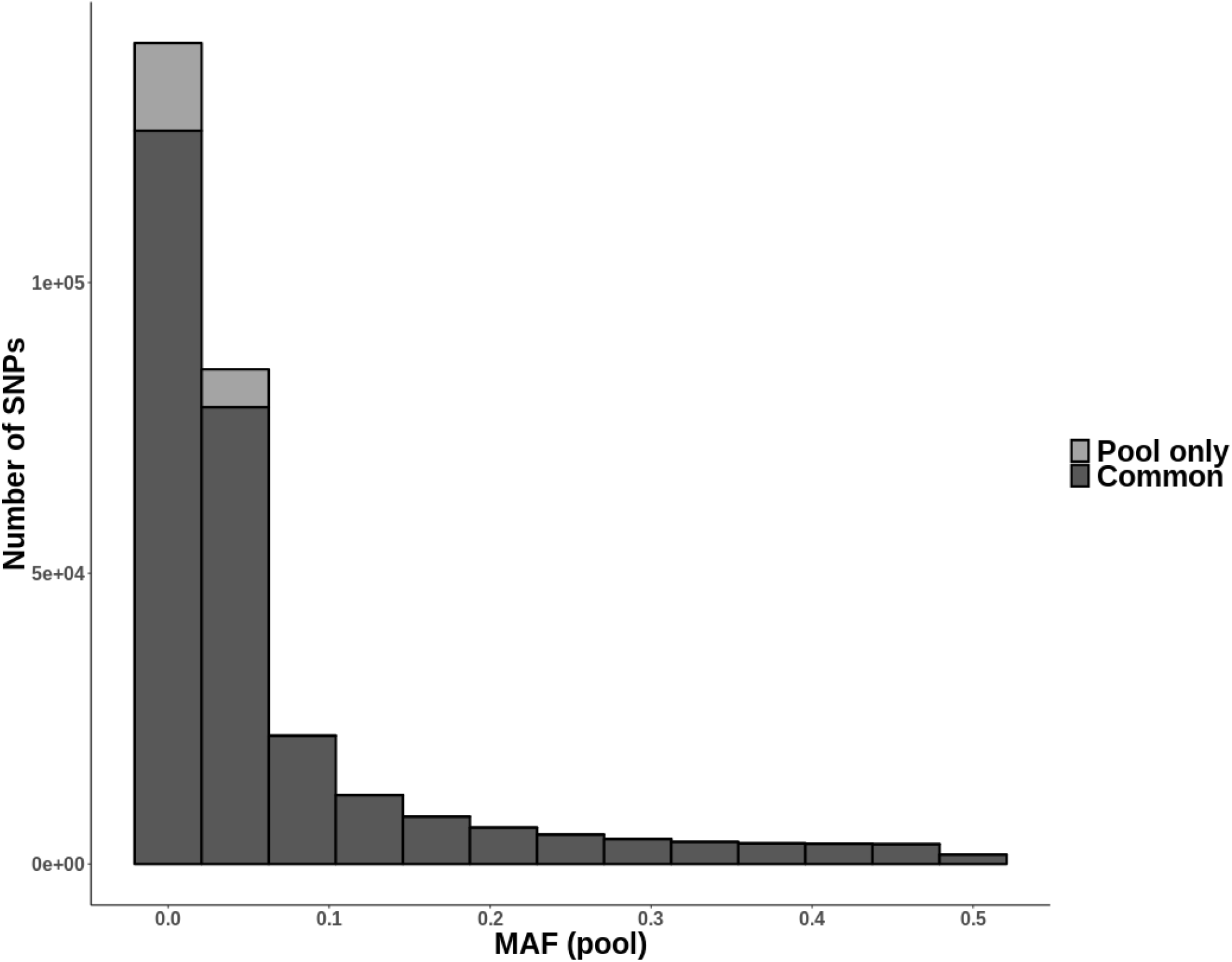
Folded-site frequency spectrum of the 300,036 filtered exonic bi-allelic SNPs (mapping onto targeted CDS sequences) according to their polymorphism in the pool and in individuals. For SNPs polymorphic in both the pool and individuals, the MAF shown is based on the pool. 86 SNPs (0.03%) that were polymorphic only in the individuals are not represented here, all have MAF<0.05.

When all 300,036 loci were considered, allele frequency estimates were strongly correlated between the pool and individuals (Pearson’s *r* = 0.99; Fig. 3A), with a good level of agreement (average bias = 9.5E10^−5^; bias SD = 0.025).

**Figure 3:**
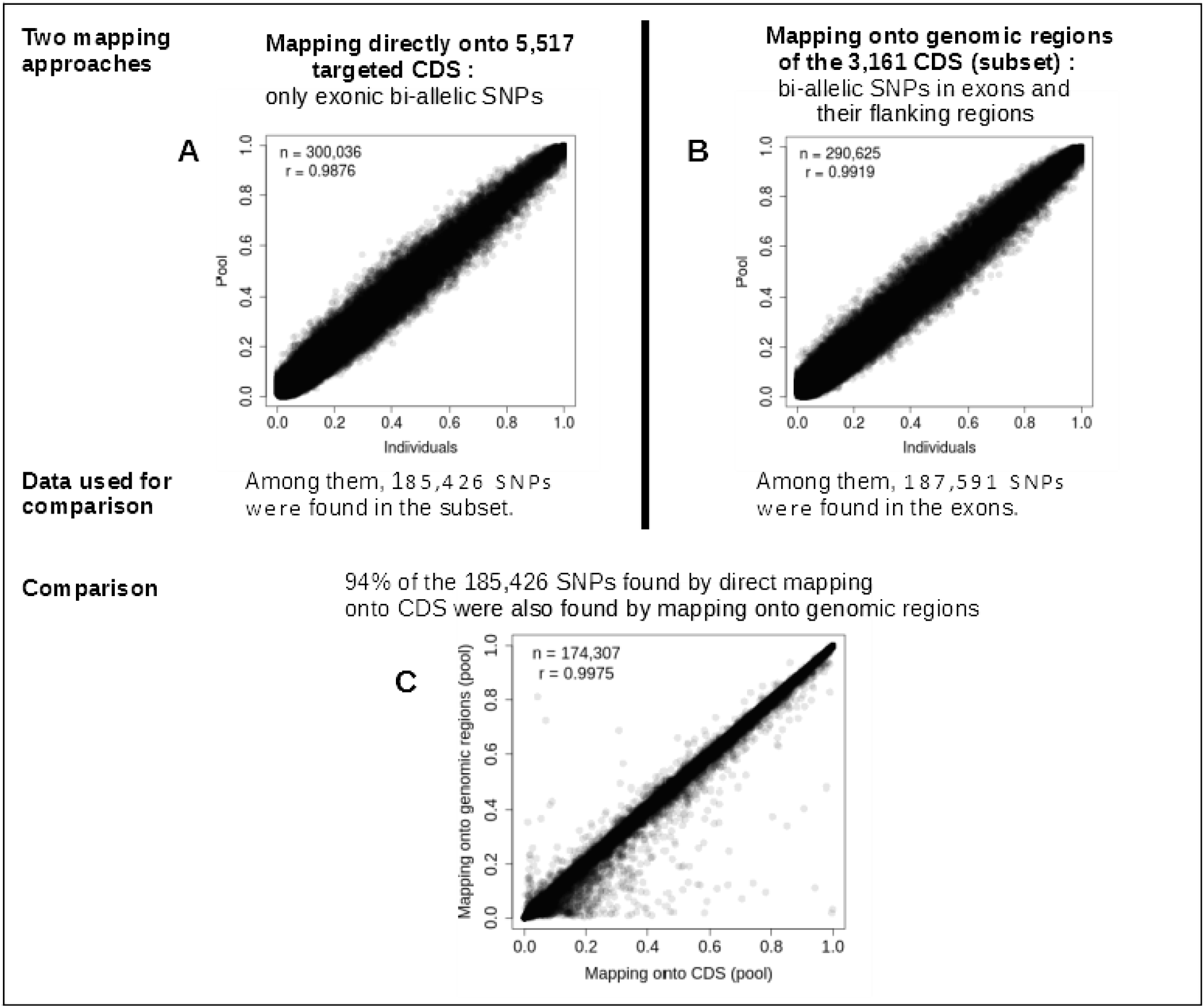
Allele frequency estimates from the population pool and individuals by two mapping approaches: **(A)** mapping reads directly onto CDS and **(B)** mapping onto genomic regions available only for a subset of target CDS. For exonic bi-allelic SNPs common to both mapping approaches, we also show **(C)** the allele frequency estimates from the pool by the two approaches. Note: Pearson correlation coefficient (*r*) are displayed.

SNP annotation revealed that 74.75% of SNPs were synonymous, whereas 24.92% were non-synonymous missense and 0.33% were non-synonymous nonsense SNPs.

### Comparison of the SNPs detected with the two mapping approaches

We evaluated the ability of the method for mapping reads directly to CDS (Fig. 1C) to call SNPs correctly by comparing the SNPs identified with this approach and those found by mapping reads to the genome (Fig. 1E). We used only the subset of 3,161 targeted CDS with a match over their entire length to sequences in the draft genome of *H. axyridis* for this comparison. For each library (pool and individuals), trimmed reads were also mapped along their entire length onto the regions of the genome corresponding to this subset (Fig. 1E). After application of the same procedure and filters for SNP calling as were used for direct mapping onto CDS sequences, 290,620 filtered bi-allelic SNPs were genotyped within the subset, including 187,591 (64.5%) in exons and 103,029 (35.5%) in the flanking regions (intron or UTR) of the exons. By mapping reads onto the genome, we demonstrated that allele frequency estimates were strongly correlated, with a good level of agreement, between the pool and individuals, for both exonic and flanking SNPs (Pearson’s *r*=0.99; average bias = −7.3E10^−4^; bias SD = 0.025; Fig. 3B).

Of the bi-allelic exonic SNPs found by direct mapping onto the targeted CDS (Fig. 3A), 185,426 mapped onto the subset of 3,161 targeted CDS, and 94% were also identified as bi-allelic SNPs by mapping to genomic regions (Fig. 1F). For these 174,307 exonic bi-allelic SNPs found with both mapping approaches (mapping onto the genome or directly onto transcripts), the allele frequency estimates obtained with the two mapping methods were highly correlated, with good level of agreement, both for the pool (Pearson’s *r*=0.998; average bias = 6.8E10^−4^; bias SD = 0.013; Fig. 3C) and for the individuals (Pearson’s *r*=0.998; average bias = 1.0E10^−3^; bias SD = 0.012).

Some of the private bi-allelic SNPs (i.e. SNPs identified in a single mapping approach) were variants initially identified with both mapping approaches but discarded during filtering in one of the approaches. For example, 1,467 bi-allelic SNPs private to the CDS mapping process had a sequencing depth above the maximum threshold, and 714 were tri-allelic SNPs when mapped onto the genome. If we exclude this latter group, 8,853 and 8,333 were found to be private to CDS mapping and mapping onto genomic regions, respectively. A non-exhaustive analysis of these positions highlighted two possible origins of private SNPs for mapping onto CDS. Some targeted regions (the full CDS or part of the CDS) have homologous copies in the genome that we were unable to identify during target selection and for which the SNPs found were not discarded with the “maximum sequencing depth” filter. We have thus identified 43 CDS (0.7%) with reads mapping onto their genomic regions in a non-unique manner, and these CDS with paralogs displayed 3,141 SNPs, all private to CDS mapping. The second origin was false SNPs occurring when the start of an intron could be aligned with the start of the next exon due to similarities between the two sequences (Fig. S2 and S6, for example). Of the SNPs private to the CDS mapping method, 6.9% (*N*=614) were positioned close to intron-exon boundary (IEB) positions (i.e. at the exact end of the exon or on the three adjacent bases), whereas only 0.6% of the 174,307 SNPs common to both mapping methods were found near IEBs.

### Prediction of intron-exon boundaries without read assembly

The method we developed for IEB detection based on the mapping of genomic reads directly onto CDS sequences (Fig. 1C) was evaluated on the 3,161 targeted CDS for which we had genomic information, and therefore knew the positions of the IEBs (Fig. 1D). An IEB signal was predicted at a particular position in the CDS if the number of reads either beginning or ending their local alignment at that position (*nbBE*) was 20 times greater (*X*=20) than the median *nbBE* value within a window of 11 bases around the position concerned, and the signal measured at that position extended to a short region of 7 bases (i.e. with a default parameter of *n*=3; see Methods section and Fig. S2). With parameters *X*=20 and *n*=3, 8,967 regions with lengths of 4 to 19 bp were predicted to be IEBs (Table 2). Of the 8,967 regions predicted to be IEBs, 8,575 actually corresponded to true IEBs (i.e. method specificity = 95.63%). Of the 8,632 regions truly containing an IEB, 8,575 were accurately predicted to be IEB regions (i.e. sensitivity = 99.34%; Table 2). Of the 862 CDS with no intron, 763 (i.e. 88.51%) were correctly predicted to have no IEBs. As expected, the use of a lower detection threshold (i.e. *X*=10) decreased the specificity of the method and increased its sensitivity (Table 2). The regions predicted to have no IEB (i.e. the - regions) were between 1 and 3,417 bp in length, with a mean length of 265.5 bp, entirely consistent with the known distribution of exon lengths (see Methods section). We counted 153 very small (<10 bp) - regions with *n=3* and only 51 with *n=5*, which attested to the possible cumulative deviation of signals of more than eight bases for a small number of IEBs. Increasing the length of the signal deviation (e.g. *n*=5 bases instead of 3) made it possible to take this situation into account in the vast majority of cases. This slightly improved performance but increased the length, and thus, the imprecision, of all predicted IEB positions (Table 2).

**Table 2:**
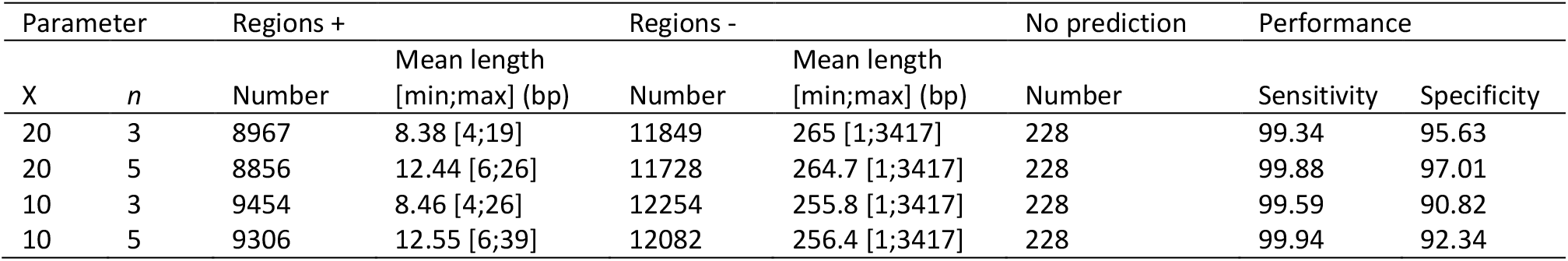
Evaluation of the IEB prediction method using pool data. The + region is a short region predicted to be an IEB. The - region is a region with no signal (i.e. predicted to be an exon). Regions not covered by reads are regions without prediction. Sensitivity is the percentage of regions with true IEBs correctly predicted to be + regions, and specificity is the percentage of + regions that are true IEBs.

| Parameter |  | Regions + |  | Regions - |  | No prediction | Performance |  |
| --- | --- | --- | --- | --- | --- | --- | --- | --- |
| X | n | Number | Mean length [min;max] (bp) | Number | Mean length [min;max] (bp) | Number | Sensitivity | Specificity |
| 20 | 3 | 8967 | 8.38 [4;19] | 11849 | 265 [1;3417] | 228 | 99.34 | 95.63 |
| 20 | 5 | 8856 | 12.44 [6;26] | 11728 | 264.7 [1;3417] | 228 | 99.88 | 97.01 |
| 10 | 3 | 9454 | 8.46 [4;26] | 12254 | 255.8 [1;3417] | 228 | 99.59 | 90.82 |
| 10 | 5 | 9306 | 12.55 [6;39] | 12082 | 256.4 [1;3417] | 228 | 99.94 | 92.34 |

## Discussion

Exome-targeted capture associated with next-generation sequencing has been mainly restricted to model species for which genomic resources are available (Ng *et al*., 2009). Transcriptome-based exome capture has proved an efficient alternative for dealing with non-model species (e.g. Bi *et al*., 2012), but is mostly used in phylogeny studies. Here, we propose an extension of this method with a high capture efficiency, providing accurate estimates of the allele frequencies of SNPs for population genomics studies. Our methodology is original because (i) it combines transcriptome-based exome capture with a pool-based approach (pooling the tissues of numerous individuals before DNA extraction), (ii) it does not require a genome sequence for the species itself or another closely related species, and (iii) it does not involve the *de novo* assembly of genomic reads. This work opens up interesting prospects for population genomics studies on non-model species because it is cheap and less labor-intensive than other methods.

### Capture efficiency

We found that capture efficiency was very high for the *H. axyridis* exome subset we targeted. The capture specificity (83.2%) achieved was much higher than reported in more classical transcriptome-based studies (range 25% - 73%), whereas capture sensitivity (93.5%) approached the highest reported values (see Puritz & Lotterhos, 2018). Moreover, despite the differences in sequencing depth observed between targets and along the length of targets, reproducibility was high between libraries, whether in terms of specificity, sensitivity or sequencing depth. Our capture approach, based on the choice of targets from the *de novo* transcriptome of the species of interest itself, without the need to identify orthologous genes in related model species, proved highly effective. This “blind” approach had only a minor impact on capture performance: we only observed a slight decrease in sequencing depth for small exons and towards the ends of the exons. Similar patterns have been observed in classical transcriptome-based exome capture in which IEBs are predicted by orthology. In both cases, this is due to the use of less efficient probes designed from the sequences of two consecutive exons and the absence of a probe straddling the end of the exon and the beginning of the intron (e.g. Bi *et al*., 2012; Neves *et al*., 2013).

The choice of targets without looking for orthology in related species was probably the cause of the slightly lower capture sensitivity in this study than in previous studies (Bi *et al*., 2012; Puritz & Lotterhos, 2018). Targets selected this way are necessarily only partly known, and some may belong to other species (e.g. symbionts or parasites) or be chimeras present in the transcriptome used. This is a small price to pay given the efficiency of the method, a feature of considerable interest for population genomics studies. The absence of searches for orthologous genes in related model species prevents the analysis from focusing purely on the most conserved genes. Moreover, our method can be applied to any species for which no closely related model species with genomic resources is available.

### SNP calling and the pool-based approach

With this method, we were able to genotype a very large number of SNPs (∼300,000) over a random subset of the exome of *H. axyridis*. The study of the subset of CDS with complete genomic sequence matches further supported the idea that direct mapping onto the CDS has probably no major impact on SNP identification or on the estimation of allele frequencies relative to mapping onto the genome. In addition to a few residual paralogous sequences inherent to any approach dealing with non-model species, we observed slightly larger numbers of SNPs called on the basis of mapping onto CDS in areas very close to the IEBs, but the proportion of SNPs involved was minimal (0.33%).

For similar sequencing efforts, strong concordance was observed between the allele frequencies for individual genotyping and those for the “pool-based” approach, whatever the mapping method used. This finding demonstrates the feasibility of this approach, in situations in which individual identification for many samples is practically impossible, technically difficult, too costly and/or unnecessary. In population genomics studies, allele frequencies are sufficient for the calculation of most summary statistics (e.g. nucleotide diversity, *F*ST, site frequency spectrum) and, thus, for addressing most of the questions posed (Schlötterer *et al*., 2014). In the specific case of exome sequencing, allele frequencies can be used for large population-scale screening of protein mutations, or to address questions related to adaptation or to the evolution of the genetic load (e.g. Gutenkunst *et al*., 2009; Yi *et al*., 2010; Casals *et al*., 2013; Popp *et al*., 2017; Castellano *et al*., 2019).

One of the limitations of pool-seq compared to individual-based approaches is that it is not optimal for studies in which analyses rely on linkage phase information. However, a prerequisite in this context is the availability of a genetic (or physical) map, which is not the case in most non-model organisms. Another weakness of pool-seq is that “populations” must be carefully defined before sequencing, as it will be difficult to detect a Wahlund effect or an unforeseen breeding system afterwards. It is therefore important to be particularly careful during the sampling step, and to be as familiar as possible with the biology of the species being studied. Finally, one more drawback of pool-seq is that it makes it impossible to remove SNPs that are not at Hardy-Weinberg equilibrium. Such disequilibrium may be caused by copy number variations or paralogous sequences resulting in false-positive SNPs and local large sequencing depth (Li, 2014). We show here that the application of our “maximum sequencing depth” filter to the pooled data made it possible to discard almost 70% of the SNPs not at equilibrium, as shown in individuals. The extremely small remaining proportion (0.34%) of SNPs not at Hardy-Weinberg equilibrium provides support for the feasibility of our pooling-based method.

### Method for predicting IEBs without read assembly

We have developed a method for detecting IEBs that is particularly useful in the case of non-model species (or species for which genome sequences are being assembled). The proposed tool (available from https://github.com/edeleury/IEB-finder) does not indicate the exact location of the IEBs, instead it identifies an area of a few bases (about eight, on average, in our case) predicted to contain an IEB. This method, which requires only the genomic reads and the target CDS sequences, was effective when evaluated on a subset of CDS for which we had a complete genomic match. Almost all the true IEBs were identified, whereas false-positives accounted for less than 5% of the predicted IEBs. Montes *et al*. (2013) proposed a similar approach, which they used to discard putative SNPs found in predicted IEB areas. They obtained better genotyping results for the selected filtered SNPs than other similar studies on non-model species. In this study, we went a step further by taking into account the possible offset of the measured signal, and by evaluating the performance of the approach on a set of transcripts for which the true intron-exon structure was known. We aimed to present evidence that this approach is promising, although further improvement is required, together with evaluation on larger dataset and at various sequencing depth levels. A method for choosing an appropriate value for parameter *X* for the correct identification of significant increases in *nbBE* depending on local sequencing depth must also be developed.

### General recommendations, conclusion and perspectives

The combination of approaches proposed here — i.e. exome transcriptome-based capture and direct mapping of reads onto targeted CDS sequences, coupled with a pool-seq approach before DNA extraction — significantly decreases costs, the laboratory time required and the complexity of the sequencing analyses required to obtain a large number of variants in exome-scale coding regions for non-model species. However, direct mapping onto target sequences has two disadvantages. First, it provides no information on flanking regions (including regulatory regions), and it is therefore not possible to genotype SNPs in these areas. Nevertheless, detecting SNP polymorphisms at exonic level would be sufficient in many cases, particularly when the in-depth analysis of a large number of synonymous and non-synonymous mutations is informative enough to address the question posed. It is still feasible to assemble the genomic reads in situations in which flanking regions appear to be of interest. Second, direct mapping onto CDS results in the detection of some false SNPs in the vicinity of IEBs. As a means of limiting this risk further, we therefore suggest using the IEB detection method proposed here, followed by the elimination of any SNPs found in the predicted IEB regions and in short regions (≤ 20 bp) surrounded by two predicted IEB regions. Using this approach on our data with the parameters *X*=20 and *n*=3, we retained a final list of 296,736 SNPs. A large proportion of the 3,300 SNPs excluded (408 SNPs, i.e. 12.36%) were not at Hardy Weinberg equilibrium in individuals, which is an additional evidence of the relevance of this filter.

The allele frequencies measured for the pool with our method were in good agreement with those obtained in the analysis of individuals. However, caution is nevertheless required with rare alleles (i.e. SNPs with low MAFs), which may be confused with sequencing errors (Bansal, 2010). We recommend the use of technical replicates. When possible, and depending on the biological question addressed, the analysis could be restricted to SNPs with MAFs above a given threshold.

In conclusion, our study demonstrates the feasibility of obtaining, at low cost, for a non-model species (i.e. without a reference genome), important and accurate information at population level concerning (i) SNP polymorphism within the exome and (ii) the general structure of the exome (intron-exon structure of the targeted genes). The results obtained with this method applied to the Asian ladybird *Harmonia axyridis* were very promising, and we now need to investigate other species with different genome sizes and complexities. Finally, the oversizing of these experiments, resulting in very high sequencing depths, suggests that it should be possible to reduce costs further by capturing multiple indexed pools in a single reaction.

## Online supplementary material

Supplementary figures and tables are available online: https://doi.org/10.1101/583534

## Data accessibility

The exome sequence capture data reported here have been deposited in the European Nucleotide Archive (accession no. PRJEB31592). The CDS sequences targeted and the description of the exon positions on the subset of transcripts have been deposited at Zenodo https://doi.org/10.5281/zenodo.2598388. The HaxR v1.0 genome sequence used in this study is available from http://bipaa.genouest.org/sp/harmonia_axyridis/ (see Gautier *et al*., 2018).

## Acknowledgements

We thank Stefan Toepfer for the *Harmonia axyridis* samples. We thank Thibaut Malausa, Arnaud Estoup, Mathieu Gautier, Fabrice Legeai, and Bernhard Gschloessl for their helpful comments, which helped us to improve this manuscript considerably. This work was funded by the INRA SPE department and the INRA UMR Institut Sophia Agrobiotech. Sequencing was performed at the GENOTOUL GeT platform (https://get.genotoul.fr/en/). Version 7 of this preprint has been peer-reviewed and recommended by Peer Community In Genomics (https://doi.org/10.24072/pci.genomics.100002).

## Author Contributions

ED, EL and TG conceived the study. AB, ED and EL prepared the samples and the libraries. ED developed the bioinformatics pipelines. ED and EL ran the analyses. ED, EL and TG wrote the paper.

