## Supplemental file for "An evaluation of pool-sequencing transcriptome-based exon capture for population genomics in non-model species"


| | |
|---|---|
| *Figure S1* | *p. 2* |
| *Figure S2* | *p. 3* |
| *Figure S3* | *p. 4* |
| *Figure S4* | *p. 5* |
| *Figure S5* | *p. 6* |
| *Figure S6* | *p. 7* |
| *Table S1* | *p. 8* |
| *Table S2* | *p. 9* |
| *Table S3* | *p. 10* |
| *Table S4* | *p. 11* |

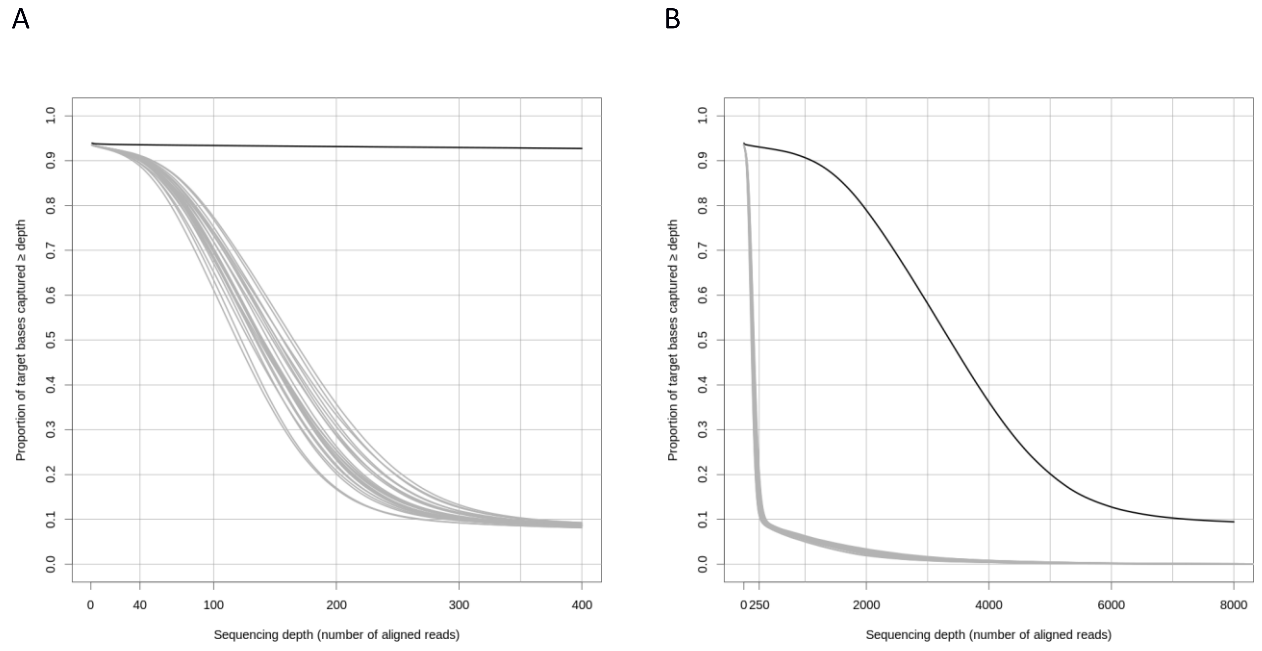


**Figure S1**: Cumulative base sequencing depth for each of the 24 genomic libraries. The libraries for the 23 individuals are represented in grey and the pool library is shown in black. A and B show exactly the same data, with different scales for the *x*-axis.

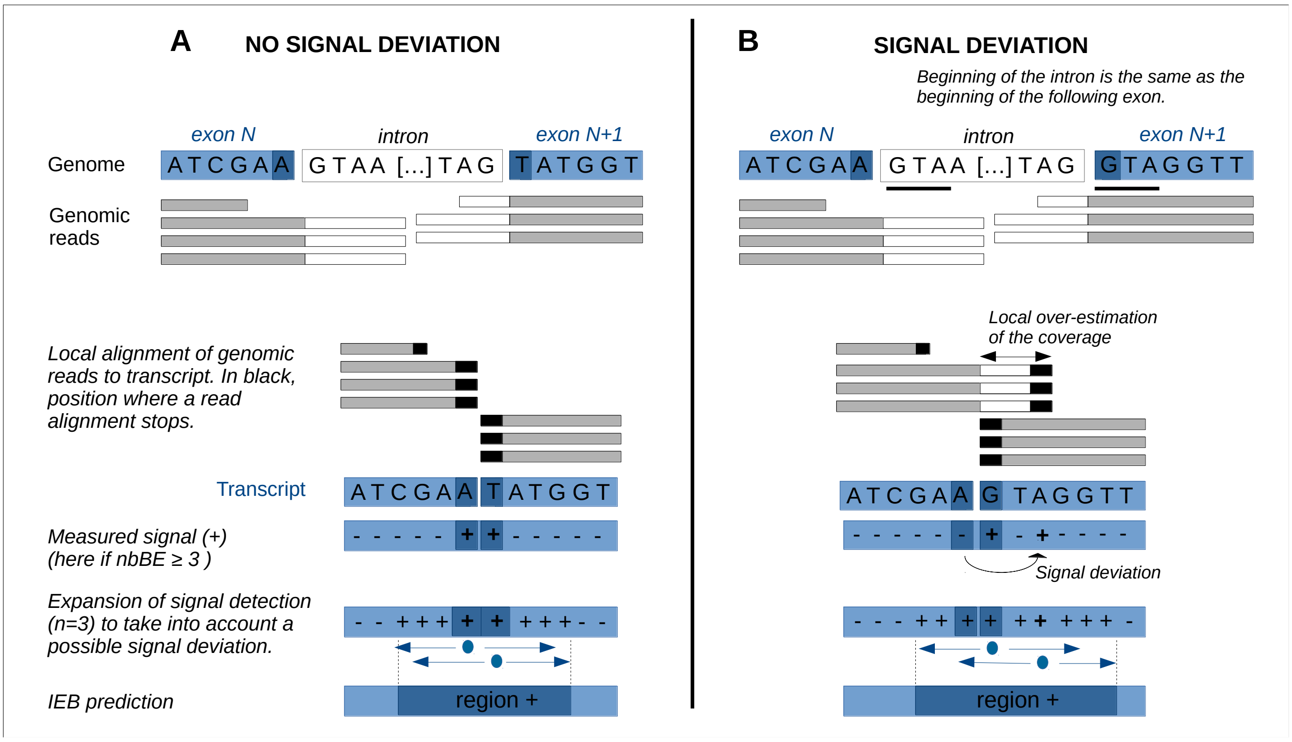


**Figure S2**: New method for predicting intron-exon boundaries (IEB) by mapping genomic reads directly onto CDS sequences and by counting, at each position, the number of reads beginning or ending at that position (nbBE). Two cases are illustrated here: on the left, when the reads stop mapping to the exact position of the exon end for the two exon ends of one IEB; on the right, when the reads stop mapping to some bases of the exon end for one of the two exon ends. The latter case is possible when the beginning of the intron is, by chance, similar to the beginning of the next exon. This can occur for both exon ends, which increases the length of the IEB prediction region (the + region).

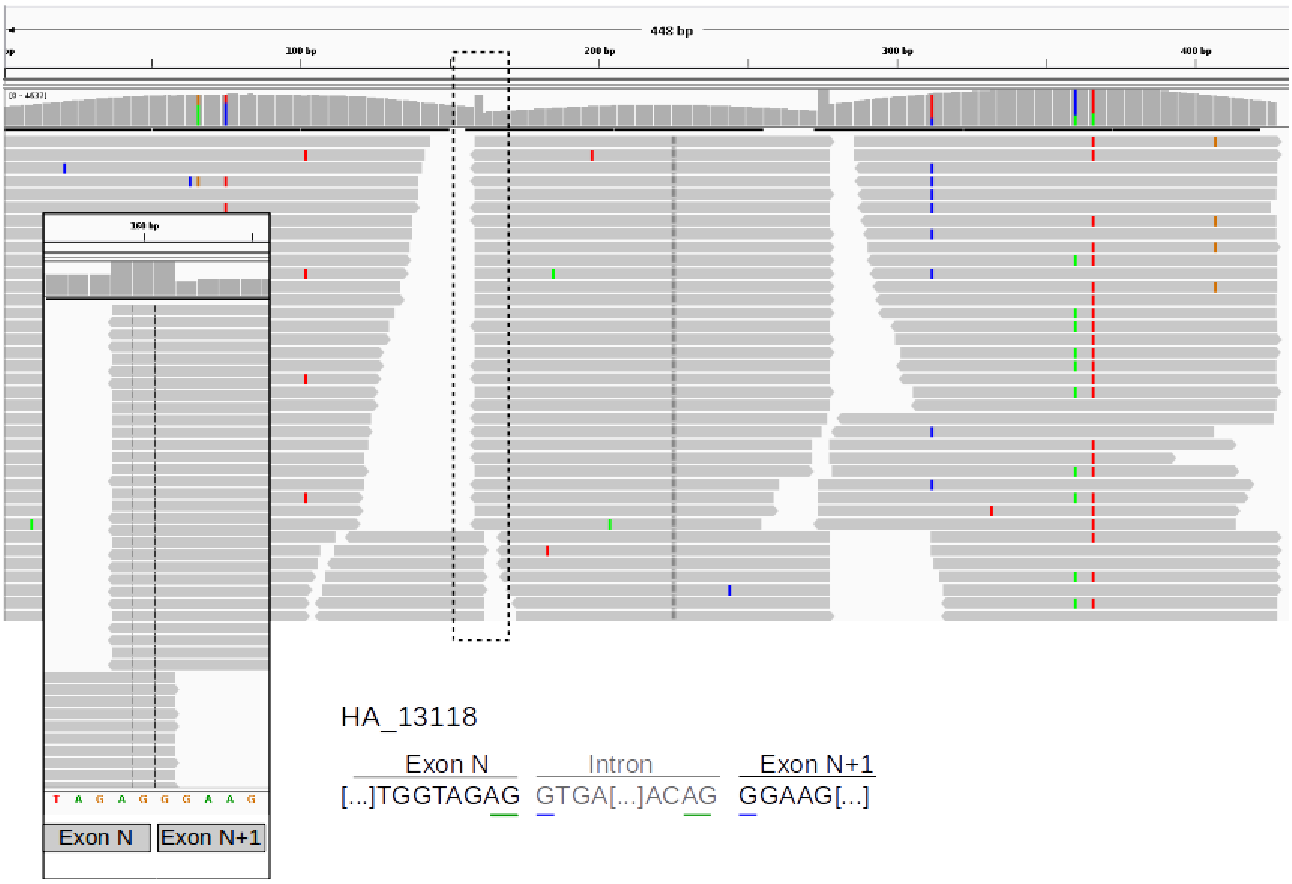


**Figure S3**: Mapping of genomic reads directly onto targeted CDS (concatenation of exons) visualized in Integrative Genomics Viewer (IGV). Reads begin or end next to an intron-exon boundary (IEB), with the possible alignment of the intron with some bases. Here, the first intron base (G) can be aligned with the first base of the exon N+1 (G), and the last two bases of the intron (AG) can be aligned with the end of exon N (AG). In total, sequencing depth is abnormally overestimated for three bases located next to this IEB (AG//G) in the CDS.

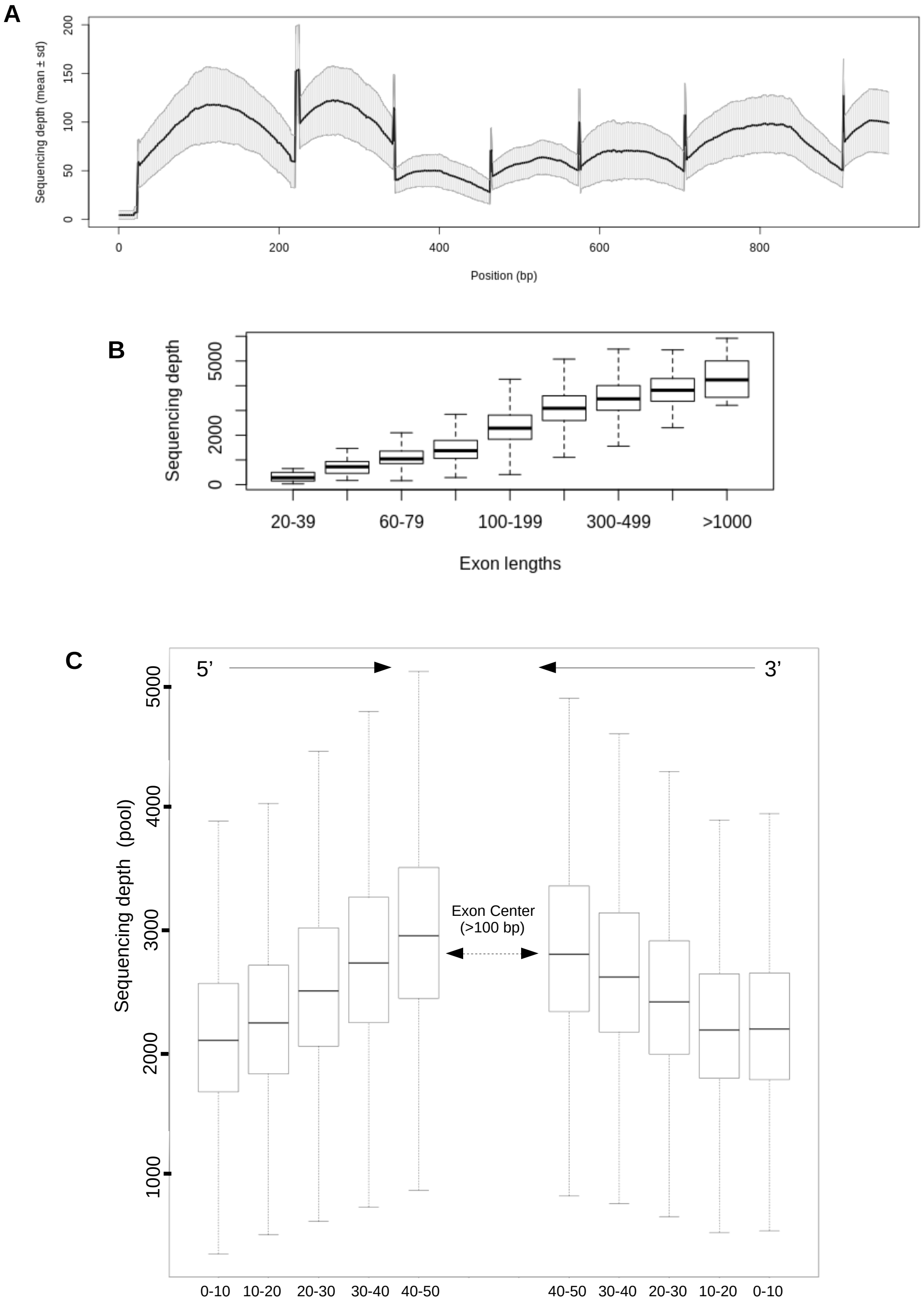


**Figure S4**: **A.** Mean sequencing depth (plus or minus standard error) for the 23 individual libraries, plotted as a function of position along the *HA_10038* coding sequence (960 bp; concatenation of 8 exons). The sequencing depth at the exon ends was overestimated, as explained in Figure S2. **B.** Mean exon sequencing depth as a function of exon length ($n$ = 6,293 complete exons ≥ 20 bp) for pool data. **C.** Mean binned sequencing depth at the ends of 200-500 bp complete exons ($n$ = 2,729 exons) for pool data. We used 10-base pair bins, with five bins for the 5' and 3' ends.

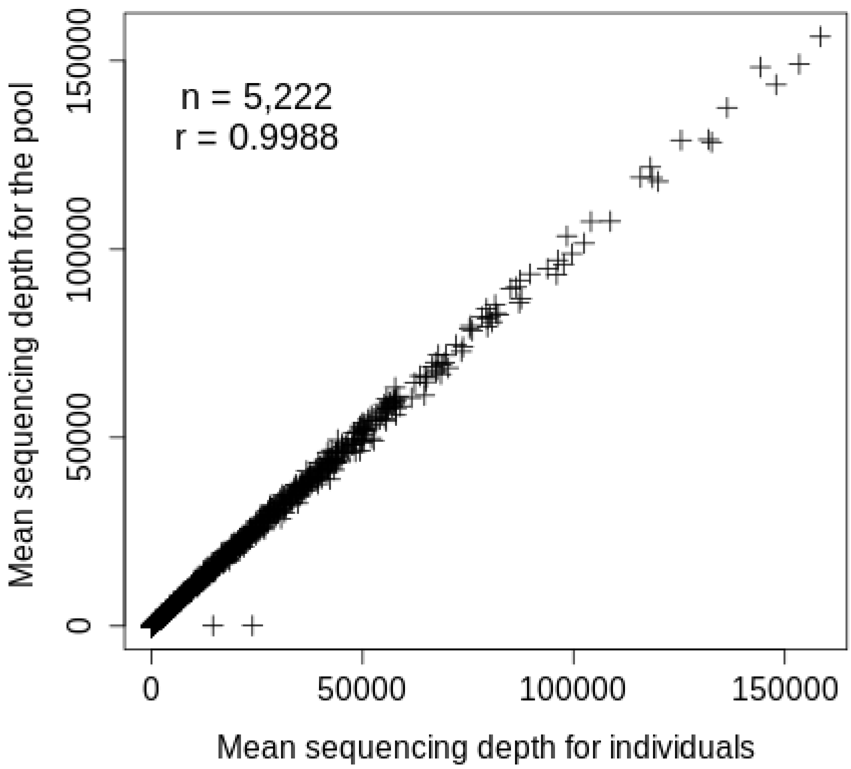


**Figure S5**:
Comparison of the mean sequencing depth of targeted CDS between the two capture methods, *i.e.* between the pool and individuals. Each cross corresponds to a CDS target. Note: 4 out-of-range CDS targets were discarded from the analysis for the purposes of representation.

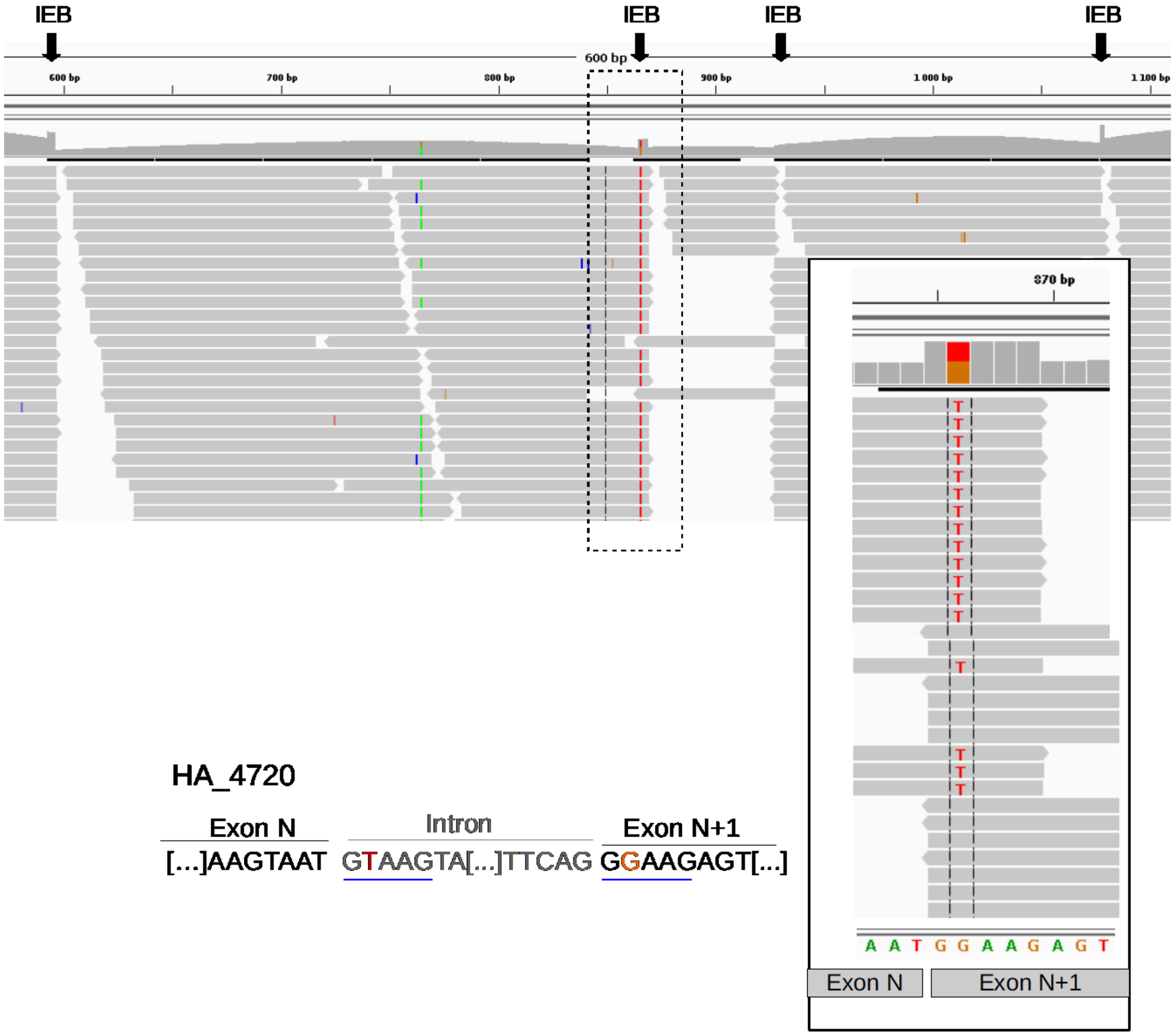


**Figure S6**: Example of a false SNP called next to an exon end (or intron-exon boundary (IEB)) when reads were mapped directly onto the target CDS. Partial IGV screenshot. Here, the beginning of the intron sequence is very similar to the first five bases of the next exon sequence in the CDS. Thus, genomic reads with the end of exon N and the start of the intron can align with the end of exon N and the start of exon N+1 over five bases. This creates an abnormally inflated estimate of sequencing depth locally for five bases at the boundary of exon N+1. An SNP is detected when the two sequences are not strictly identical, as in this example.

| Library Name | Number of read pairs | Mbases | Proportion of reads (%) | Number of cleaned read pairs | Number of cleaned reads (SE1 only) | Number of cleaned reads (SE2 only) | Total number of cleaned reads | Proportion of read retained (%) |
|---|---|---|---|---|---|---|---|---|
| Pool of 36 non-indexed individuals | 186998614 | 56100 | 50,2 | 156104105 | 26592490 | 1700304 | 340501004 | 91,0 |
| 23 indexed individuals (subtotal) | 185457659 | 55637 | 49,8 | 156088966 | 23928203 | 2421740 | 338527875 | 91,3 |
| *Individual 1* | *7241970* | *2173* | *1,94* | *6029132* | *1045507* | *70861* | *13174632* | *91,0* |
| *Individual 2* | *7484575* | *2245* | *2,01* | *6202580* | *1099934* | *74390* | *13579484* | *90,7* |
| *Individual 3* | *8732346* | *2620* | *2,34* | *7237004* | *1300459* | *83530* | *15857997* | *90,8* |
| *Individual 4* | *7668116* | *2300* | *2,06* | *6353240* | *1138260* | *76576* | *13921316* | *90,8* |
| *Individual 5* | *7738008* | *2321* | *2,08* | *6420474* | *1139591* | *77887* | *14058426* | *90,8* |
| *Individual 6* | *7637178* | *2291* | *2,05* | *6332647* | *1122654* | *77082* | *13865030* | *90,8* |
| *Individual 7* | *8391493* | *2517* | *2,25* | *6943476* | *1256004* | *81538* | *15224494* | *90,7* |
| *Individual 8* | *6912561* | *2074* | *1,86* | *5474368* | *1285154* | *64563* | *12298453* | *89,0* |
| *Individual 9* | *7811654* | *2343* | *2,10* | *6564169* | *903887* | *147683* | *14179908* | *90,8* |
| *Individual 10* | *9149586* | *2745* | *2,46* | *7932068* | *972713* | *95969* | *16932818* | *92,5* |
| *Individual 11* | *8531966* | *2560* | *2,29* | *7074679* | *1060746* | *155945* | *15366049* | *90,0* |
| *Individual 12* | *7875656* | *2363* | *2,11* | *6306581* | *747908* | *508170* | *13869240* | *88,1* |
| *Individual 13* | *7974834* | *2392* | *2,14* | *6902758* | *871979* | *79481* | *14756976* | *92,5* |
| *Individual 14* | *7762438* | *2329* | *2,08* | *6636308* | *933444* | *79203* | *14285263* | *92,0* |
| *Individual 15* | *7504558* | *2251* | *2,01* | *6469430* | *839092* | *77207* | *13855159* | *92,3* |
| *Individual 16* | *8987541* | *2696* | *2,41* | *7771107* | *1002259* | *91165* | *16635638* | *92,5* |
| *Individual 17* | *8071329* | *2421* | *2,17* | *6904584* | *966861* | *75809* | *14851838* | *92,0* |
| *Individual 18* | *8371274* | *2511* | *2,25* | *7028527* | *1140522* | *86402* | *15283978* | *91,3* |
| *Individual 19* | *7629129* | *2289* | *2,05* | *6561116* | *877072* | *78807* | *14078111* | *92,3* |
| *Individual 20* | *7726175* | *2318* | *2,07* | *6526912* | *1014177* | *77100* | *14145101* | *91,5* |
| *Individual 21* | *8405897* | *2522* | *2,26* | *7174677* | *1045475* | *78480* | *15473309* | *92,0* |
| *Individual 22* | *9176292* | *2753* | *2,46* | *7805586* | *1145586* | *90937* | *16847695* | *91,8* |
| *Individual 23* | *8673083* | *2602* | *2,33* | *7437543* | *1018919* | *92955* | *15986960* | *92,2* |
| ***Total*** | ***372456273*** | ***111737*** | ***100*** | ***312193071*** | ***50520693*** | ***4122044*** | ***679028879*** | ***91,2*** |

**Table S1**: Summary statistics for the raw sequence data obtained for the 24 libraries

| Library Name | Total number of cleaned reads | Mb (cleaned reads) | unmapped reads | % | Reads aligned not properly or not on only one target | % | Reads aligned properly (*) on only one target | % | Number of mapped read pairs | % | Orphelan mapped reads | % | Number of Mb of mapped read that can be aligned | % | median read alignment length (pb) | mean read alignment length (pb) | min | max | % of reads that mapped on their entire length |
|---|---|---|---|---|---|---|---|---|---|---|---|---|---|---|---|---|---|---|---|
| Pool of non-indexed 36 individuals | 340501004 | 47879 | 43807484 | 12.9 | 11961756 | 3.5 | 284731764 | 83.6 | 124044982 | 77.2 | 36641801 | 22.8 | 35926 | 75.0 | 144 | 126.2 | 21 | 194 | 62.7 |
| 23 indexed individuals (subtotal) | 338527875 | 47564 | 47298784 | 14.0 | 11459979 | 3.4 | 280467456 | 82.8 | 122895052 | 78.4 | 33938762 | 21.6 | 35315 | 74.2 | 144 | 126.3 | 21 | 194 | 62.7 |
| *Individual 1* | *13174632* | *1857* | *1727954* | *13.1* | *389241* | *3.0* | *11755781* | *89.2* | *4857144* | *78.8* | *1302903* | *21.2* | *1397.7* | *75.3* | *144* | *126.9* | *21* | *190* | *62.3* |
| *Individual 2* | *13579484* | *1912* | *1912219* | *14.1* | *442293* | *3.3* | *11224972* | *82.7* | *4932128* | *78.4* | *1360716* | *21.6* | *1419.9* | *74.3* | *144* | *126.6* | *21* | *184* | *62.2* |
| *Individual 3* | *15857997* | *2234* | *2060717* | *13.0* | *513572* | *3.2* | *13283708* | *83.8* | *5852549* | *78.8* | *1578610* | *21.2* | *1685.0* | *75.4* | *144* | *126.9* | *21* | *184* | *62.2* |
| *Individual 4* | *13921316* | *1963* | *1817460* | *13.1* | *446801* | *3.2* | *11657055* | *83.7* | *5148113* | *79.1* | *1360829* | *20.9* | *1477.6* | *75.3* | *144* | *126.8* | *21* | *181* | *61.5* |
| *Individual 5* | *14058426* | *1983* | *1842578* | *13.1* | *460067* | *3.3* | *11755781* | *83.6* | *5182201* | *78.8* | *1391379* | *21.2* | *1490.6* | *75.2* | *144* | *126.9* | *21* | *183* | *61.8* |
| *Individual 6* | *13865030* | *1952* | *1784540* | *12.9* | *445871* | *3.2* | *11634619* | *83.9* | *5120592* | *78.6* | *1393435* | *21.4* | *1473.9* | *75.5* | *144* | *126.8* | *21* | *181* | *61.9* |
| *Individual 7* | *15224494* | *2146* | *2064586* | *13.6* | *486293* | *3.2* | *12673615* | *83.2* | *5581393* | *78.7* | *1510829* | *21.3* | *1607.3* | *74.9* | *144* | *126.9* | *21* | *185* | *62.0* |
| *Individual 8* | *12298453* | *1730* | *1585884* | *12.9* | *371706* | *3.0* | *10340863* | *84.1* | *4472482* | *76.2* | *1395899* | *23.8* | *1310.1* | *75.7* | *144* | *126.8* | *21* | *185* | *60.9* |
| *Individual 9* | *14179908* | *1956* | *2072547* | *14.6* | *471677* | *3.3* | *11635684* | *82.1* | *5091130* | *77.8* | *1453424* | *22.2* | *1444.8* | *73.9* | *143* | *124.2* | *21* | *190* | *63.6* |
| *Individual 10* | *16932818* | *2382* | *2545433* | *15.0* | *627331* | *3.7* | *13760054* | *81.3* | *6045511* | *78.4* | *1669032* | *21.6* | *1733.7* | *72.8* | *144* | *126.1* | *21* | *194* | *63.2* |
| *Individual 11* | *15366049* | *2107* | *2449822* | *15.9* | *531803* | *3.5* | *12384424* | *80.6* | *5270579* | *74.1* | *1843266* | *25.9* | *1530.3* | *72.6* | *143* | *123.6* | *21* | *180* | *64.3* |
| *Individual 12* | *13869240* | *1922* | *2064938* | *14.9* | *474583* | *3.4* | *11329719* | *81.7* | *4843138* | *74.7* | *1643443* | *25.3* | *1413.5* | *73.6* | *144* | *124.8* | *21* | *186* | *64.0* |
| *Individual 13* | *14756976* | *2079* | *2127230* | *14.4* | *526246* | *3.6* | *12103500* | *82.0* | *5337625* | *78.9* | *1428250* | *21.1* | *1529.5* | *73.6* | *144* | *126.4* | *21* | *183* | *63.3* |
| *Individual 14* | *14285263* | *2013* | *1975586* | *13.8* | *504009* | *3.5* | *11805668* | *82.6* | *5189804* | *78.4* | *1426060* | *21.6* | *1492.8* | *74.2* | *144* | *126.5* | *21* | *190* | *63.1* |
| *Individual 15* | *13855159* | *1950* | *1973303* | *14.2* | *504201* | *3.6* | *11377655* | *82.1* | *5002107* | *78.5* | *1373441* | *21.5* | *1435.4* | *73.6* | *144* | *126.2* | *21* | *181* | *63.2* |
| *Individual 16* | *16635638* | *2348* | *2346627* | *14.1* | *581732* | *3.5* | *13707279* | *82.4* | *6091118* | *80.0* | *1525043* | *20.0* | *1739.6* | *74.1* | *144* | *127.0* | *21* | *186* | *63.5* |
| *Individual 17* | *14851838* | *2087* | *2136572* | *14.4* | *518154* | *3.5* | *12197112* | *82.1* | *5377092* | *78.8* | *1442928* | *21.2* | *1538.4* | *73.7* | *144* | *126.2* | *21* | *187* | *63.0* |
| *Individual 18* | *15283978* | *2154* | *2081198* | *13.6* | *494861* | *3.2* | *12707919* | *83.1* | *5618944* | *79.3* | *1470031* | *20.7* | *1610.5* | *74.8* | *144* | *126.8* | *21* | *183* | *62.1* |
| *Individual 19* | *14078111* | *1984* | *1994794* | *14.2* | *503180* | *3.6* | *11580137* | *82.3* | *5108075* | *78.9* | *1363987* | *21.1* | *1462.5* | *73.7* | *144* | *126.4* | *21* | *182* | *62.8* |
| *Individual 20* | *14145101* | *1993* | *1959427* | *13.9* | *479115* | *3.4* | *11706559* | *82.8* | *5167455* | *79.0* | *1371649* | *21.0* | *1479.8* | *74.3* | *144* | *126.5* | *21* | *183* | *62.2* |
| *Individual 21* | *15473309* | *2183* | *2142096* | *13.8* | *527369* | *3.4* | *12803844* | *82.7* | *5672944* | *79.6* | *1457956* | *20.4* | *1621.9* | *74.3* | *144* | *126.7* | *21* | *188* | *62.6* |
| *Individual 22* | *16847695* | *2375* | *2377652* | *14.1* | *610065* | *3.6* | *13859978* | *82.3* | *6093890* | *78.5* | *1672198* | *21.5* | *1751.8* | *73.8* | *144* | *126.5* | *21* | *188* | *62.8* |
| *Individual 23* | *15986960* | *2256* | *2255621* | *14.1* | *549809* | *3.4* | *13181530* | *82.5* | *5839038* | *79.5* | *1503454* | *20.5* | *1669.0* | *74.0* | *144* | *126.7* | *21* | *187* | *62.8* |
| ***Total*** | ***679028879*** | ***95444*** | ***91106268*** | ***13.4*** | ***23421735*** | ***3.4*** | ***565199220*** | ***83.2*** | ***246940034*** | ***77.8*** | ***70580563*** | ***22.2*** | ***71242*** | ***74.6*** | ***144*** | ***126.3*** | | | ***62.7*** |

**Table S2**: Raw mapping data for all 24 libraries

| Library Name | TARGET BASE LEVEL<br>Nb bases with no read | Nb of bases with at least one read | % | Mean base sequencing depth | Median base sequencing depth | Number of base with sequencing depth <300X and <7000X for individuals and pool respectively | % | GLOBAL TARGET LEVEL<br>Nb targets with no read | Number of targets with at least one read | % | Mean target sequencing depth | Median target sequencing depth |
|---|---|---|---|---|---|---|---|---|---|---|---|---|
| Pool of non-indexed 36 individuals | 326294 | 5021167 | *93.9* | 6634.8 | 3361 | 4795633 | *89.7* | 500 | 5217 | *91.3* | 7033.0 | 3324 |
| *Individual 1* | 351465 | 4995996 | *93.4* | 239.3 | 119 | 4854127 | *90.8* | 545 | 5172 | *90.5* | 254.9 | 119 |
| *Individual 2* | 348242 | 4999219 | *93.5* | 262.2 | 131 | 4827714 | *90.3* | 537 | 5180 | *90.6* | 278.9 | 130 |
| *Individual 3* | 346493 | 5000968 | *93.5* | 311.2 | 152 | 4709553 | *88.1* | 540 | 5177 | *90.6* | 331.2 | 154 |
| *Individual 4* | 347151 | 5000310 | *93.5* | 272.9 | 139 | 4801695 | *89.8* | 542 | 5175 | *90.5* | 291.2 | 139 |
| *Individual 5* | 347769 | 4999692 | *93.5* | 275.2 | 140 | 4788898 | *89.6* | 544 | 5173 | *90.5* | 292.8 | 140 |
| *Individual 6* | 349565 | 4997896 | *93.5* | 272.1 | 137 | 4786866 | *89.5* | 545 | 5172 | *90.5* | 291.6 | 140 |
| *Individual 7* | 346194 | 5001267 | *93.5* | 296.7 | 149 | 4745769 | *88.7* | 543 | 5174 | *90.5* | 315.5 | 149 |
| *Individual 8* | 348816 | 4998645 | *93.5* | 241.9 | 122 | 4852703 | *90.7* | 545 | 5172 | *90.5* | 257.6 | 122 |
| *Individual 9* | 347880 | 4999581 | *93.5* | 266.8 | 135 | 4819917 | *90.1* | 540 | 5177 | *90.6* | 280.6 | 134 |
| *Individual 10* | 348145 | 4999316 | *93.5* | 320.0 | 162 | 4671124 | *87.4* | 536 | 5181 | *90.6* | 338.9 | 160 |
| *Individual 11* | 345535 | 5001926 | *93.5* | 282.5 | 139 | 4782468 | *89.4* | 548 | 5169 | *90.4* | 295.9 | 141 |
| *Individual 12* | 347692 | 4999769 | *93.5* | 260.9 | 128 | 4823433 | *90.2* | 544 | 5173 | *90.5* | 275.6 | 130 |
| *Individual 13* | 345963 | 5001498 | *93.5* | 282.4 | 140 | 4785941 | *89.5* | 533 | 5184 | *90.7* | 300.9 | 142 |
| *Individual 14* | 348232 | 4999229 | *93.5* | 275.6 | 138 | 4807067 | *89.9* | 544 | 5173 | *90.5* | 291.9 | 138 |
| *Individual 15* | 348292 | 4999169 | *93.5* | 264.9 | 134 | 4813076 | *90.0* | 544 | 5173 | *90.5* | 280.8 | 133 |
| *Individual 16* | 347384 | 5000077 | *93.5* | 321.2 | 157 | 4662713 | *87.2* | 543 | 5174 | *90.5* | 341.2 | 158 |
| *Individual 17* | 347555 | 4999906 | *93.5* | 284.0 | 144 | 4771824 | *89.2* | 541 | 5176 | *90.5* | 300.4 | 144 |
| *Individual 18* | 350460 | 4997001 | *93.4* | 297.4 | 148 | 4730760 | *88.5* | 547 | 5170 | *90.4* | 317.4 | 150 |
| *Individual 19* | 348109 | 4999352 | *93.5* | 269.9 | 138 | 4802299 | *89.8* | 547 | 5170 | *90.4* | 285.9 | 137 |
| *Individual 20* | 348670 | 4998791 | *93.5* | 273.2 | 139 | 4801834 | *89.8* | 543 | 5174 | *90.5* | 289.9 | 139 |
| *Individual 21* | 346829 | 5000632 | *93.5* | 299.5 | 151 | 4741120 | *88.7* | 546 | 5171 | *90.4* | 317.0 | 151 |
| *Individual 22* | 347987 | 4999474 | *93.5* | 323.3 | 165 | 4638896 | *86.7* | 543 | 5174 | *90.5* | 343.7 | 165 |
| *Individual 23* | 349634 | 4997827 | *93.5* | 308.1 | 156 | 4713310 | *88.1* | 544 | 5173 | *90.5* | 328.3 | 156 |
| **Considering all 24 libraries** | **323418** | **5024043** | ***94.0*** | **13136** | **6662** | **4796032** | ***89.7*** | **491** | **5226** | ***91.4*** | **12738.3** | **6331.0** |

**Table S3**: Raw results for capture efficiency per target for all 24 libraries

| Library Name | Correctly aligned reads | Number of duplicates | % |
|---|---|---|---|
| Pool of 36 non-indexed individuals | 284731764 | 273926275 | 96,2 |
| 23 indexed individuals (subtotal) | 279728866 | 185312226 | 66,2 |
| *Individual 1* | *11017191* | *7141232* | *64,8* |
| *Individual 2* | *11224972* | *7285322* | *64,9* |
| *Individual 3* | *13283708* | *9043890* | *68,1* |
| *Individual 4* | *11657055* | *7562879* | *64,9* |
| *Individual 5* | *11755781* | *7705899* | *65,5* |
| *Individual 6* | *11634619* | *7581803* | *65,2* |
| *Individual 7* | *12673615* | *8488778* | *67,0* |
| *Individual 8* | *10340863* | *6572290* | *63,6* |
| *Individual 9* | *11635684* | *7569467* | *65,1* |
| *Individual 10* | *13760054* | *9338775* | *67,9* |
| *Individual 11* | *12384424* | *8214520* | *66,3* |
| *Individual 12* | *11329719* | *7347433* | *64,9* |
| *Individual 13* | *12103500* | *7972917* | *65,9* |
| *Individual 14* | *11805668* | *7760721* | *65,7* |
| *Individual 15* | *11377655* | *7384171* | *64,9* |
| *Individual 16* | *13707279* | *9442127* | *68,9* |
| *Individual 17* | *12197112* | *8058067* | *66,1* |
| *Individual 18* | *12707919* | *8588109* | *67,6* |
| *Individual 19* | *11580137* | *7506713* | *64,8* |
| *Individual 20* | *11706559* | *7684259* | *65,6* |
| *Individual 21* | *12803844* | *8592530* | *67,1* |
| *Individual 22* | *13859978* | *9530119* | *68,8* |
| *Individual 23* | *13181530* | *8940205* | *67,8* |

**Table S4**: Number of reads scored as duplicates for the 24 libraries.
